# Gene and cell line efficiency of CRISPR computed by tensor decomposition in genome-wide CRISPR-Cas9 knockout screens

**DOI:** 10.1101/2025.06.12.659265

**Authors:** Y-H Taguchi, Turki Turki

**Affiliations:** Department of Physics, Chuo University, 1-13-27 Kasuga, Bunkyo-ku, Tokyo, 112-8551, Japan; Department of Computer Science, King Abdulaziz University, Jeddah, 21589, Saudi Arabia

**Keywords:** unsupervised learning, tensor decomposition, CRISPR, sgRNA, genome wide analysis, novel AI application in CRISPR-Cas9

## Abstract

Genome-wide CRISPR-Cas9 knockout screens are often used to experimentally evaluate gene function. However, the efficacy of individual sgRNAs targeting unique genes varies and is difficult to integrate. In this study, tensor decomposition (TD) was used to integrate multiple sgRNAs and sgRNA profiles simultaneously. Thus, TD can discriminate between essential and non-essential genes with the performance comparative to that of Joint analysis of CRISPR/-Cas9 knockout screens (JACKS), a type of SOTA that previously outperformed various other SOTA. In addition, although TD uses simple linear algebra, it can achieve good performance even without control samples, without which JACKS cannot be performed. Moreover, because raw and logarithmic values can achieve similar performances through TD for the largest dataset among the tested datasets, taking logarithmic values as has been done frequently, which is questioned. In conclusion, TD is the first method that can integrate multiple sgRNAs attributed to single a target and sgRNA profiles at the beginning simultaneously and can achieve a performance comparable to that of JACKS.

## 1 Introduction

Genome-wide CRISPR-Cas9 knockout screens are useful methods to evaluate the function of individual genes [1]. In contrast to gene expression analysis, using which we can only estimate the activity of individual genes indirectly because the amount of expression does not always reflect the functionality of genes, genome-wide CRISPR-Cas9 knockout screens can measure the functionality of genes more directly because the functions of the knocked out genes are missing. However, because the efficacy of individual sgRNAs is not stable, multiple sgRNAs targeting individual genes must be prepared. Subsequently, the outputs from the individual sgRNAs must be integrated. The integration of multiple sgRNAs into a single gene is problematic because it affects the final results.

Various sophisticated methods have been proposed to address these problems. Model-based analysis of genome-wide CRISPR-Cas9 knockout (MAGeCK) computes gene-level scores while assuming that the outputs can obey negative signed binomial distribution [2]. MAGeCK has been improved and can learn using the ENCODE dataset (MAGeCK maximum likelihood estimation (MLE) [3]. Bayesian analysis of gene essentiality lethality (BAGEL) is a Bayesian classifier that learns essential and non-essential genes as references and scores the necessity of individual genes. BAGEL specializes in identifying essential genes through negative selection [4]. Cancer essentiality analysis by RNAi and CRISPR screens (CERES) can correct the effects of copy number variation [5] and score the dependency [6] of individual genes by integrating multiple sgRNAs [7]. Score tracking of sgRNA enrichment (STARS) can score genes based on the most condensed genes and is typically used for positive selection [8]. Joint analysis of CRISPR/Cas9 knockout screens (JACKS) can integrate screening data, including multiple samples and conditions. and more precisely evaluate the activity and necessity of individual sgRNAs [9].

Besides these CRISPR specific methods, tensor decomposition (TD) can integrate multiple outputs [10], regardless to what the multiple outputs are. TD can integrate not only gene expression, but also various other features, including DNA methylation, histone modification, miRNAs, and protein–protein interactions. Thus, not surprisingly, TD can successfully integrate the outputs from multiple sgRNAs that target a unique gene. In addition to the integration of multiple sgRNAs, TD can integrate multiple sgRNA profiles at the same time, whereas none of the above-mentioned methods can do this; individual profiles must be analyzed separately one by one, and only their outcomes can be integrated.

To determine whether TD can successfully integrate the outputs from multiple sgRNAs that target a unique gene as well as multiple sgRNA profiles simultaneously, we applied TD to multiple datasets of sgRNA profiles for which multiple methods have been tested [9] in a study in which JACKS was proposed. Thus, TD successfully discriminates essential genes from non-essential genes similar to JACKS, which previously outperformed various conventional methods [9]; however, TD is forced to analyze datasets without control samples, without which other methods cannot be performed. This is a significant advantage of the TD over various conventional methods that use complicated advanced modeling.

## 2 Results

Figure 1 presents an outline of the analysis flow.

**Fig. 1.**
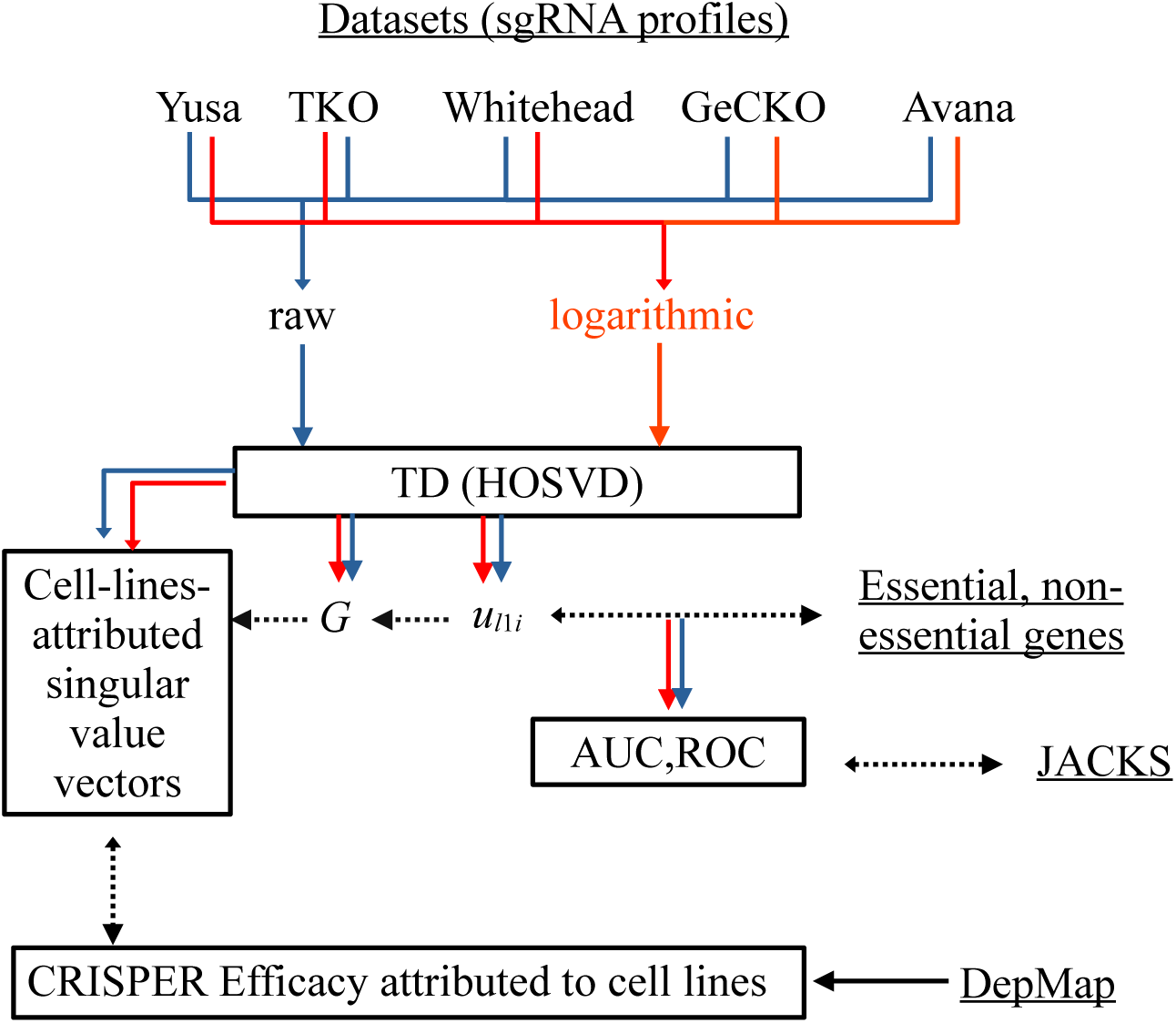
Analysis flow used in this study. Starting from five datasets comprising sgRNA profiles, the profiles are formatted as tensors with raw and logarithmic values. TD is applied to tensors, which gives *G* (core tensors) as well as singular value vectors attributed to cell lines, genes, and sgRNAs. Singular value vectors attributed to genes (denoted as *u_ℓ_*_1_ *_i_*) are evaluated by discriminating between essential and non-essential genes. Their performances are compared with those of JACKS. The cell-line-attributed singular value vectors are selected based on singular value vectors attributed to genes through *G*s. The selected cell-line-attributed singular value vectors are evaluated, and some of them are further compared with the CRISPR efficacy attributed to cell lines from DepMap databases. Blue and red lines show analysis flows for raw and logarithmic values, respectively. Dotted arrows show the comparisons to evaluate the performances of TD. Data sources of the underlined terms are from outside ones.

### 2.1 AUC, ROC, and comparison with JACKS

First, we attempted to identify which singular value vector attributed to genes, *u_ℓ_*_1 *i*_, could discriminate essential genes from nonessential genes to determine the best area under the curve (AUC) when discriminating essential genes from non-essential genes.

Table 1 lists the results. The performances in Table 1 are generally good because the AUCs for most models are as large as 0.8. To compare these performances with those of previous methods, we compared our results with those of JACKS, which outperformed numerous methods proposed earlier (Table 2) [9]. Clearly, the performances were comparable. In addition, a comparison of the receiver operating characteristic (ROC) curves revealed that JACKS and TD not only achieve similar AUCs (Tables 1 and 2) but also share similar ROC curve shapes (Fig. 2). Because the same AUC value does not always mean that the ROC curve shapes will be similar, the fact that TD and JACKS share similar ROC curve shapes means that TD and JACKS work through similar mechanisms. This is remarkable because TD employs simple linear algebra, whereas JACKS employs advanced and sophisticated Bayesian methods. Furthermore, TD does not always include control samples, without which JACKS cannot perform for some of five datasets (i.e., GeCKOv2 and Avana).

**Fig. 2.**
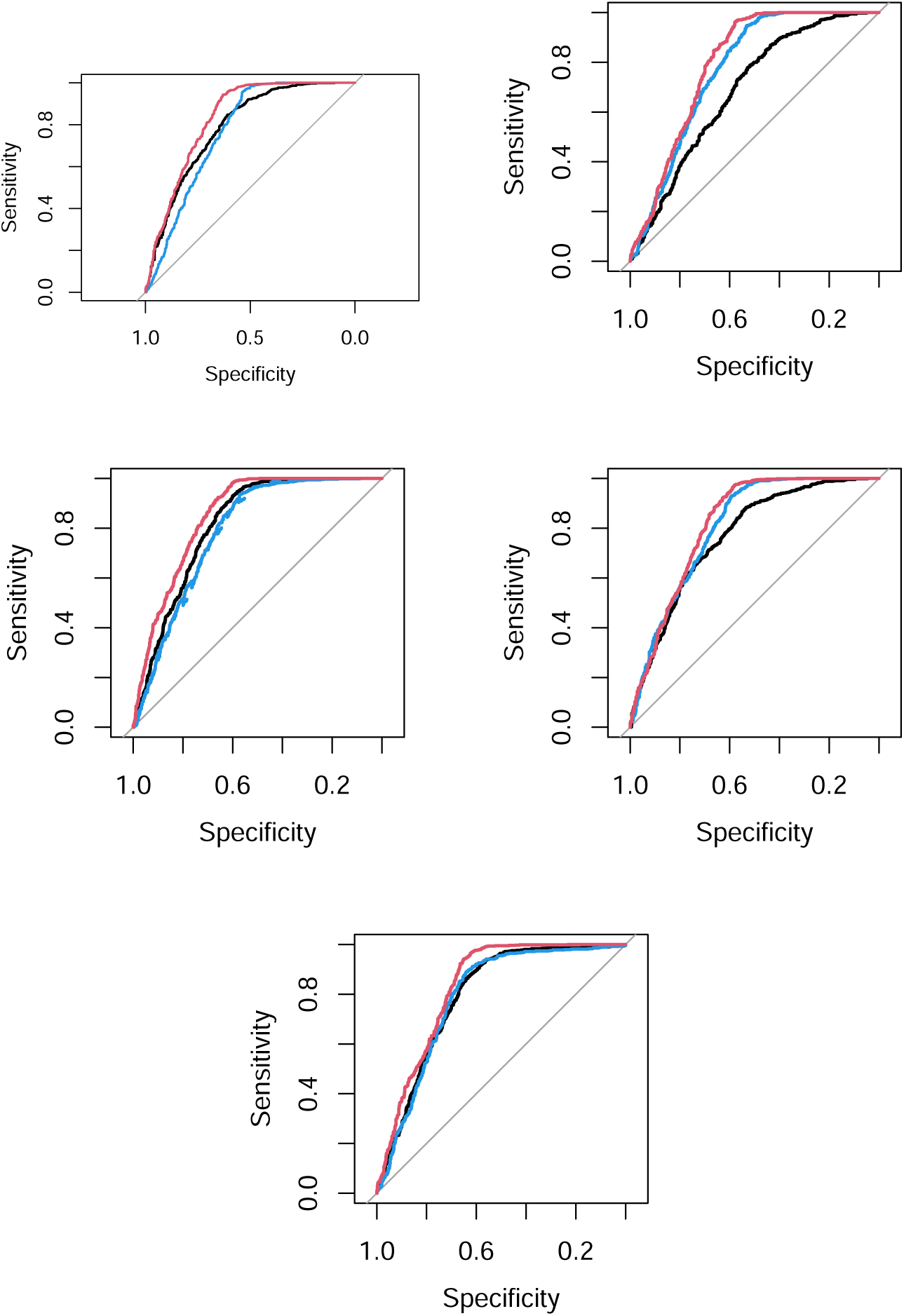
ROCs for the discrimination between essential (positive set) and non-essential genes (negative set) by TD and JACKS. The corresponding AUC values are listed in Tables 1 and 2. Black: TD with raw value, blue: TD with logarithmic values, red: JACKS. Top left: Yusa V1.0, top right: TKOv1, mid left: Whitehead, mid right: GeCKOv2, bottom Avana.

**Table 1.** The “best” singular value vectors attributed to genes, *u_ℓ_*_1_ *i*, that can discriminate essential genes from non-essential genes. For whitehead datasets, because there are two singular value vectors attributed to genes that have similarly good performances, we included both in the table. The standard errors caused by random selection of sgRNA are always less than 0.01.

| Cell lines | raw or logarithmic | $\ell_1$ | AUC |
| --- | --- | --- | --- |
| Yusa V1.0 | raw | 7 | 0.79 |
|  | logarithmic | 6 | 0.77 |
| TKv1 | raw | 1 | 0.68 |
|  | logarithmic | 5 | 0.77 |
| Whitehead | raw | 11 | 0.81 |
|  | logarithmic | 2 | 0.79 |
|  | logarithmic | 12 | 0.78 |
| GeCKOv2 | raw | 5 | 0.77 |
|  | logarithmic | 5 | 0.81 |
| Avana | raw | 5 | 0.79 |
|  | logarithmic | 5 | 0.79 |

**Table 2.** AUC to differentiate between essential (positive set) and non-essential genes (negative set) by JACKS. JACKS standard refers to AUC computed by following standard procedure. DepMap corresponds to Chronos.

| cell lines | Yusa V1.0 | TKv1 | Whitehead | GeCKOv2 | Avana |
| --- | --- | --- | --- | --- | --- |
| JACKS | 0.83 | 0.82 | 0.85 | 0.79 | 0.83 |
| JACKS standard | 0.81 | 0.75 | 0.83 | 0.81 | 0.81 |
| DepMap | — | — | — | — | 0.84 |

Thus, TD can achieve comparative performance to one of the SOTA methods, despite employing simple linear algebra and without using controls.

### 2.2 Singular value vectors attributed to cell lines

Although TD successfully achieved comparative performance to that of JACKS, a type of SOTA method, several aspects remain for discussion. First, we must clarify which singular value vectors attributed to the gene perform the best. As shown in Table 1, *ℓ*_1_ values associated with the best performance fluctuate significantly. For example, the 12th singular value vector attributed to genes, *u*_12*i*_, can achieve an almost similar AUC value with the first one in the Whitehead dataset. In general, as *ℓ*_1_ increases, its contribution to the original tensor decreases. Thus, a larger *ℓ*_1_ is less likely to have biological significance because of its noisy nature. Why can such a large *ℓ*_1_ have biological meaning for the Whitehead dataset?

To interpret the singular value vectors attributed to genes, which singular value vectors attributed to the cell lines are related to the selected singular value vectors attributed to the genes must be determined (Table 3). For this purpose, it is easier to determine whether *G*(*ℓ*_1_*ℓ*_2_*ℓ*_3_) or *G*(*ℓ*_1_*ℓ*_2_*ℓ*_3_*ℓ*_4_) have larger absolute values with fixed *ℓ*_1_. Subsequently, by inspecting the singular value vectors attributed to the cell lines with the specified *G*, we can determine why a specific *ℓ*_1_ is associated with the best performance. In the following subsections, we discuss this point individually for each of the five datasets because the interpretation is not common.

**Table 3.**
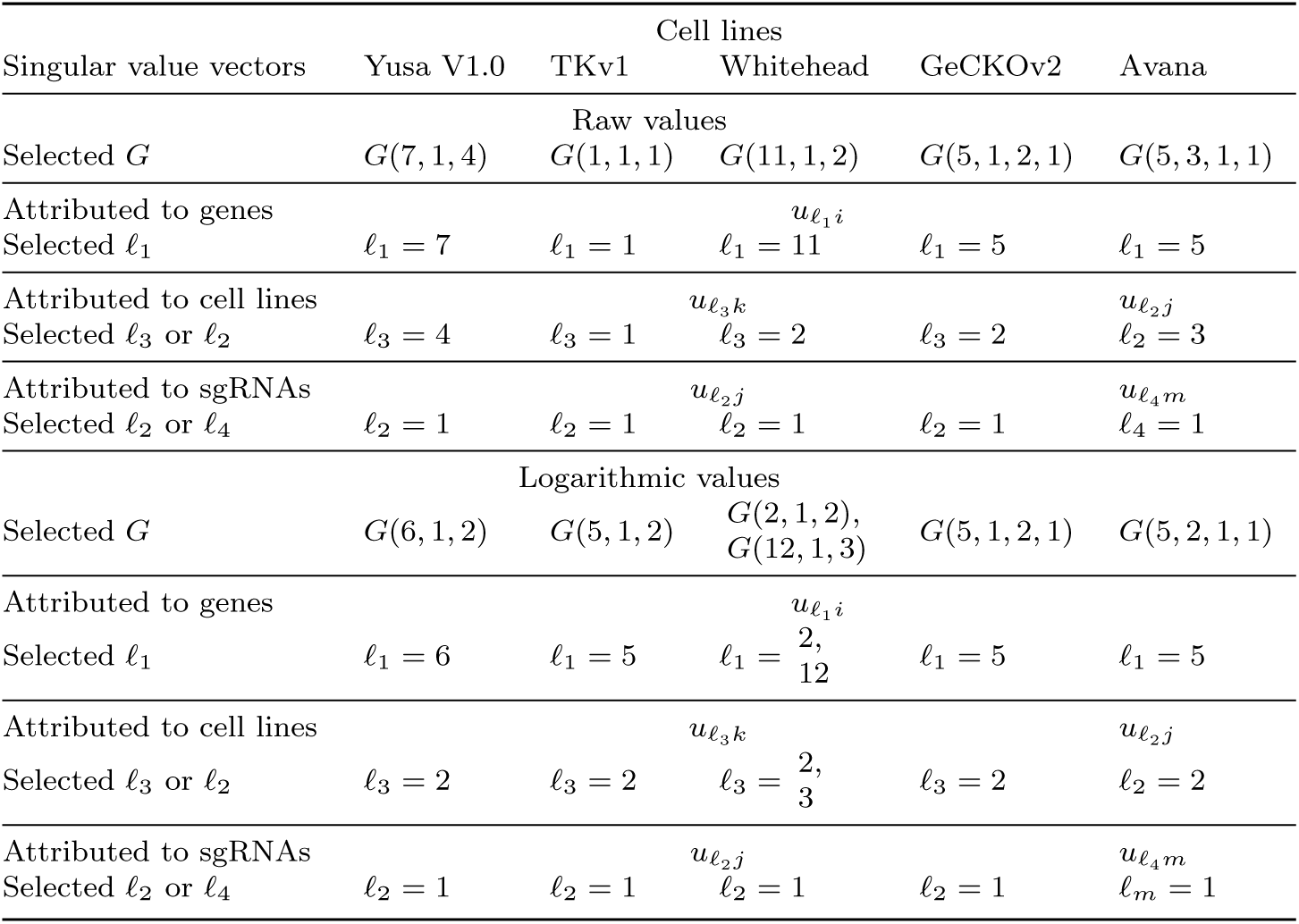
Summary of the selected singular value vectors. First, *ℓ*_1_s (attributed to genes) are selected based on the AUC values (Table 1). Subsequently, *G*s with the largest absolute values are selected (Figs. 3, 4, 6, 8, and 9). *ℓ*_2_, *ℓ*_3_, and *ℓ*_4_ (attributed to genes and sgRNA) are selected based on the selected *G*s.

#### 2.2.1 Yusa V1.0

First, we considered Yusa V1.0. The upper two panels in Figure 3 show the absolute values of *G* in descending order as well as a boxplot of *u_ℓ_*_3_ *_k_* s identified based on the *G* values. For raw and logarithmic values, *G*(7, 1, 4) and *G*(6, 1, 2) have the largest absolute values with fixed *ℓ*_1_ = 7 and *ℓ*_1_ = 6, respectively. When investigating the corresponding singular value vectors attributed to cell lines, *u*_4*k*_ for raw values and *u*_2*k*_ for logarithmic values, we know that the control profiles differ from those of the others if we observe the two lower panels in Fig. 3 (no other singular value vectors attributed to cell lines exhibit the separation of controls from other cell lines as large as these ones, not shown here). Thus, some specific singular value vectors attributed to genes are associated with the largest AUC because they are associated with cell-line-attributed singular value vectors that have the largest separation between controls and others.

**Fig. 3.**
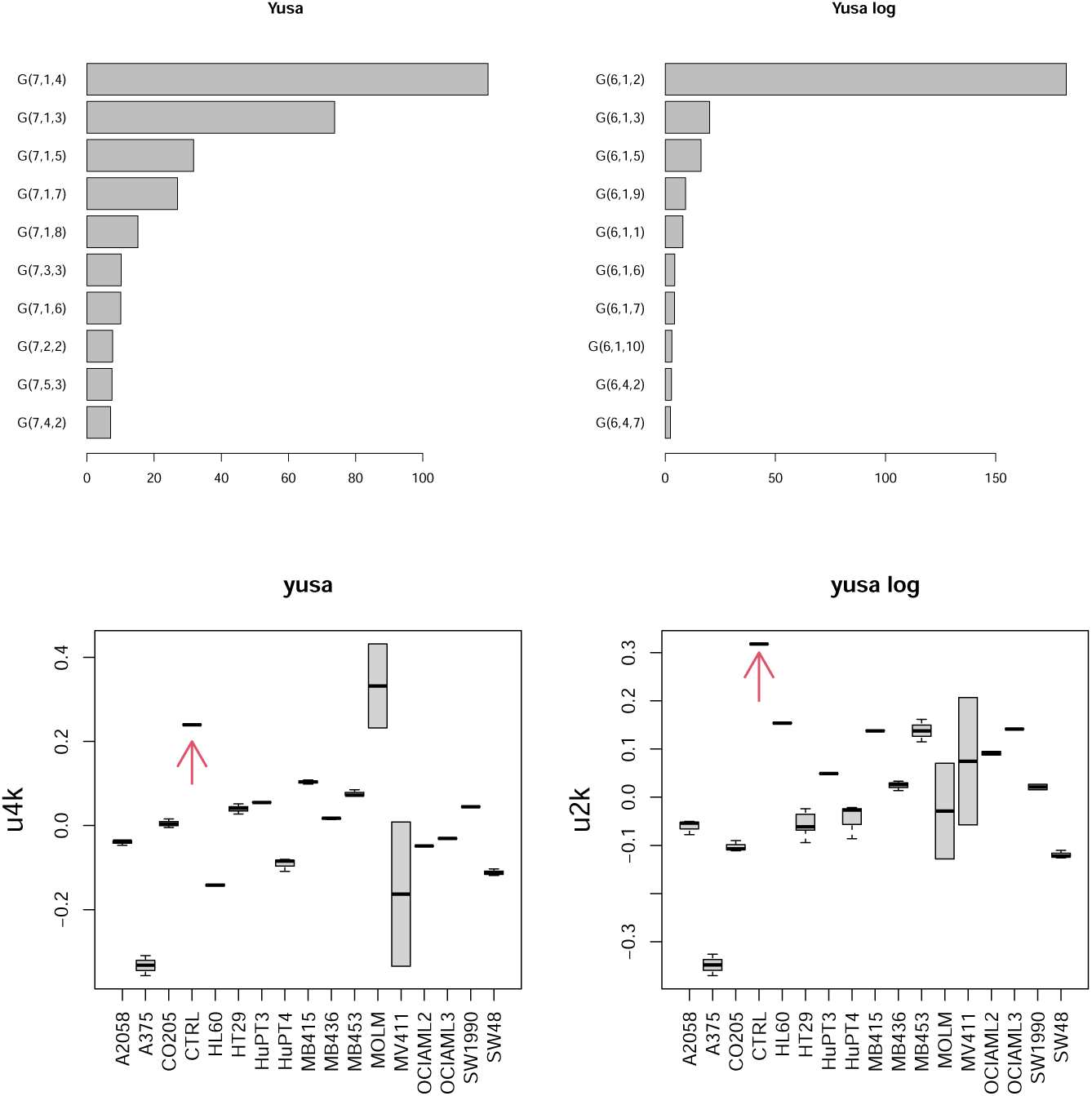
Upper row: The absolute values of *G*(*ℓ*_1_*ℓ*_2_*ℓ*_3_) in the descending order for the Yusa V1.0 dataset. *ℓ*_3_ = 4 and *ℓ*_3_ = 2 are selected for raw (left) and logarithmic vales (right), respectively. Lower raw: The singular value vectors attributed to cell lines and associated with the largest absolute *G* value with a fixed *ℓ*_1_. Left: *u*_4*k*_ for raw values and right: *u*_3*k*_ for logarithmic values. Red arrows denote controls.

#### 2.2.2 TKOv1

Similar observations were made for other datasets. For TKOv1, *G*(1, 1, 1) for raw values with *ℓ*_1_ = 1 and *G*(5, 1, 2) for logarithmic values with *ℓ*_1_ = 5 have the largest absolute values, respectively (Fig. 4). This implies that *u*_1*k*_ is expected to be associated with the selected *u*_1*i*_ for raw values, whereas *u*_2*k*_ is expected to be associated with the selected *u*_5*i*_. Thus, we should expect that *u*_1*k*_ for raw values and *u*_2*k*_ for logarithmic values have a clear separation between T0 (in some sense, it is regarded as a control because T0 is at the time point where the treatment is performed) and the others. In contrast to this expectation, no *u_ℓ_*_3_ *_k_* s attributed to cell lines exhibited a clear separation between T0 and the others for raw values, whereas *u*_2*k*_ attributed to cell lines exhibited a clear separation between T0 and the others for logarithmic values, as expected (Fig. 5). This may coincide with the observation that the raw values for the TKOv1 dataset had the lowest AUC, 0.68 (Table 1). This means that the success of TD depends on whether we can occasionally obtain cell line-attributed singular value vectors associated with the separation between the control and others.

**Fig. 4.**
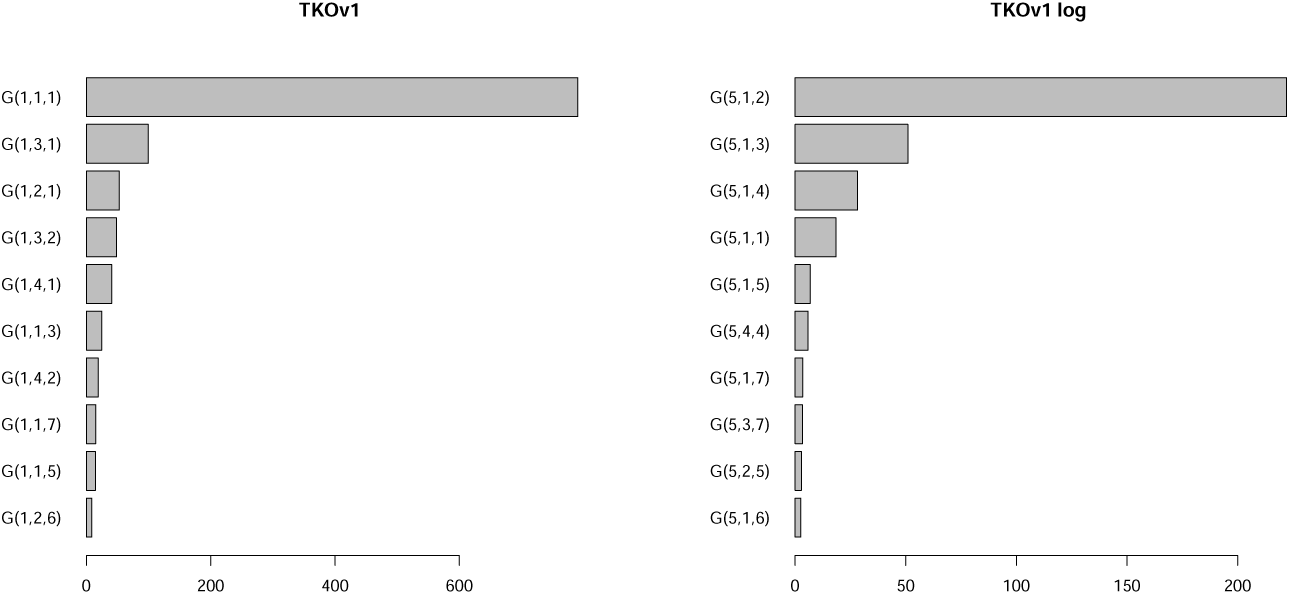
The absolute values of *G*(*ℓ*_1_*ℓ*_2_*ℓ*_3_) in descending order for the TKOv1 dataset. *ℓ*_1_ = 1 and *ℓ*_1_ = 5 are selected for raw (left) and logarithmic vales (right), respectively.

**Fig. 5.**
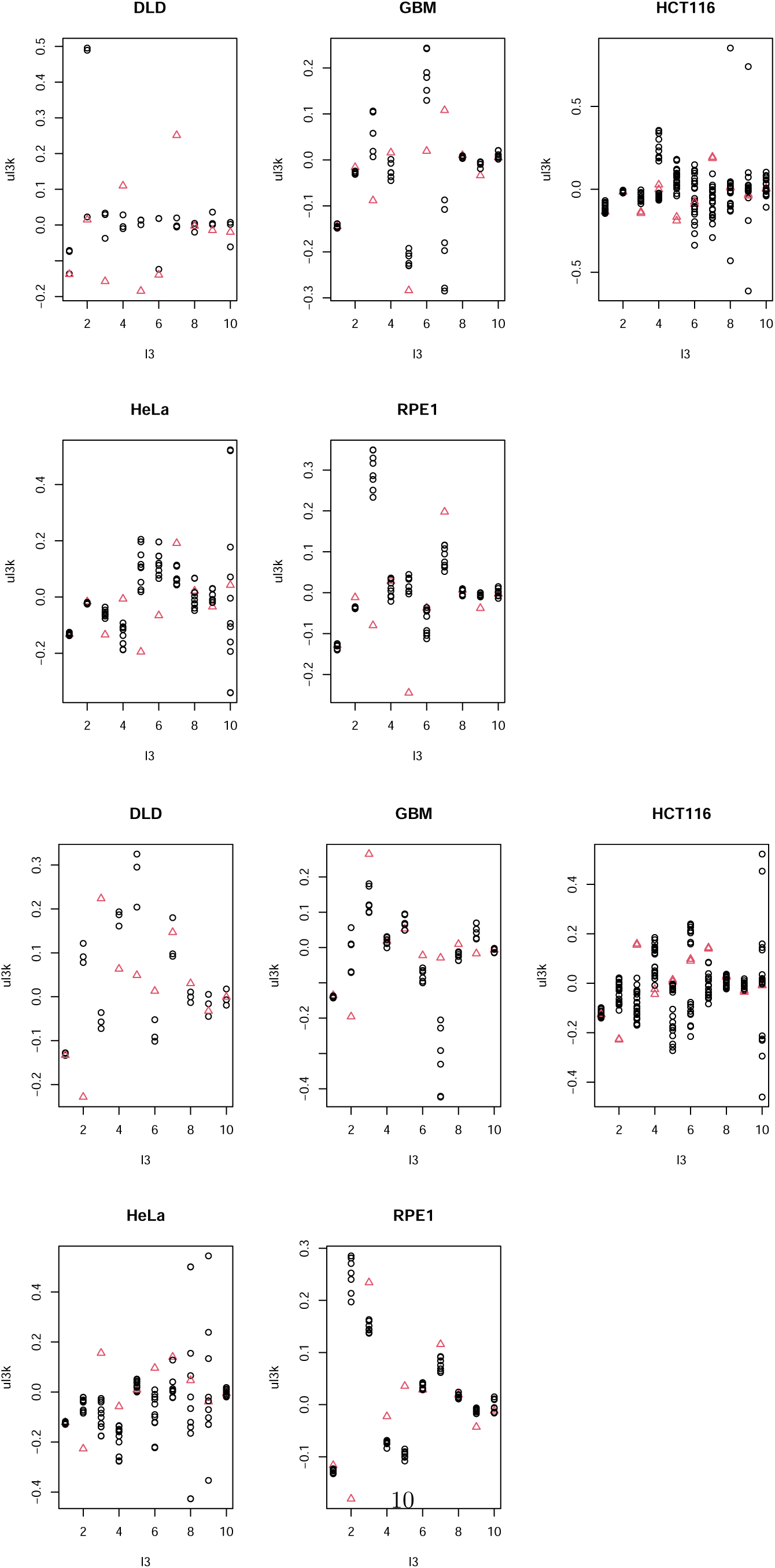
The singular value vectors attributed to cell lines, *u_ℓ_*_3_ *_k_* , for the five cell lines in the TKOv1 datasets. The first and second rows: for raw values; the third and fourth rows: for logarithmic values. Red triangles: T0; black circles: others.

#### 2.2.3 Whitehead

The whitehead dataset yields similar results. These datasets are composed of pairs of initial and final stages. *G*(11, 1, 2) for raw values with *ℓ*_1_ = 1, *G*(2, 1, 2) for logarithmic values with *ℓ*_1_ = 2, and *G*(12, 1, 3) for logarithmic values with *ℓ*_1_ = 12 having the largest absolute values (Fig. 6). Thus, *ℓ*_3_ = 2 is selected for the raw values, and *ℓ*_3_ = 2, 3 is selected for the logarithmic values. As can be seen in Fig. 7, *u*_2*k*_ for raw values and *u*_2*k*_ as well as *u*_3*k*_ for logarithmic values are associated with a large separation between the initial and final values. As the initial can be regarded as a control in some sense, a large AUC is available when singular value vectors attributed to genes are associated with cell-line-attributed singular value vectors associated with a large separation between the control and others.

**Fig. 6.**
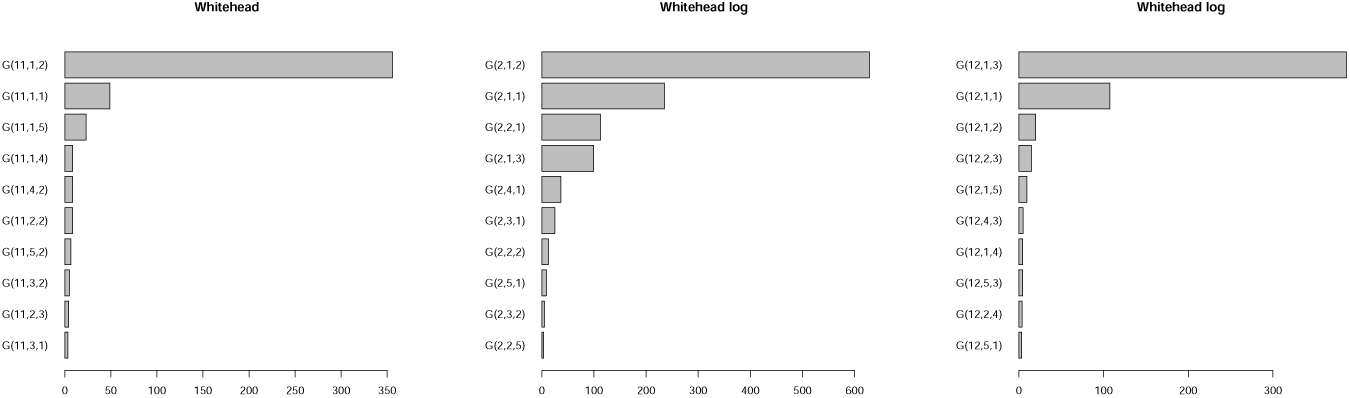
The absolute values of *G*(*ℓ*_1_*ℓ*_2_*ℓ*_3_) in descending order for the whitehead dataset. Left: for raw values with *ℓ*_1_ = 11 (*ℓ*_3_ = 1 is selected); middle: for logarithmic values with *ℓ*_1_ = 2 (*ℓ*_3_ = 2 is selected); right: for logarithmic values with *ℓ*_1_ = 12 (*ℓ*_3_ = 3 is selected).

**Fig. 7.**
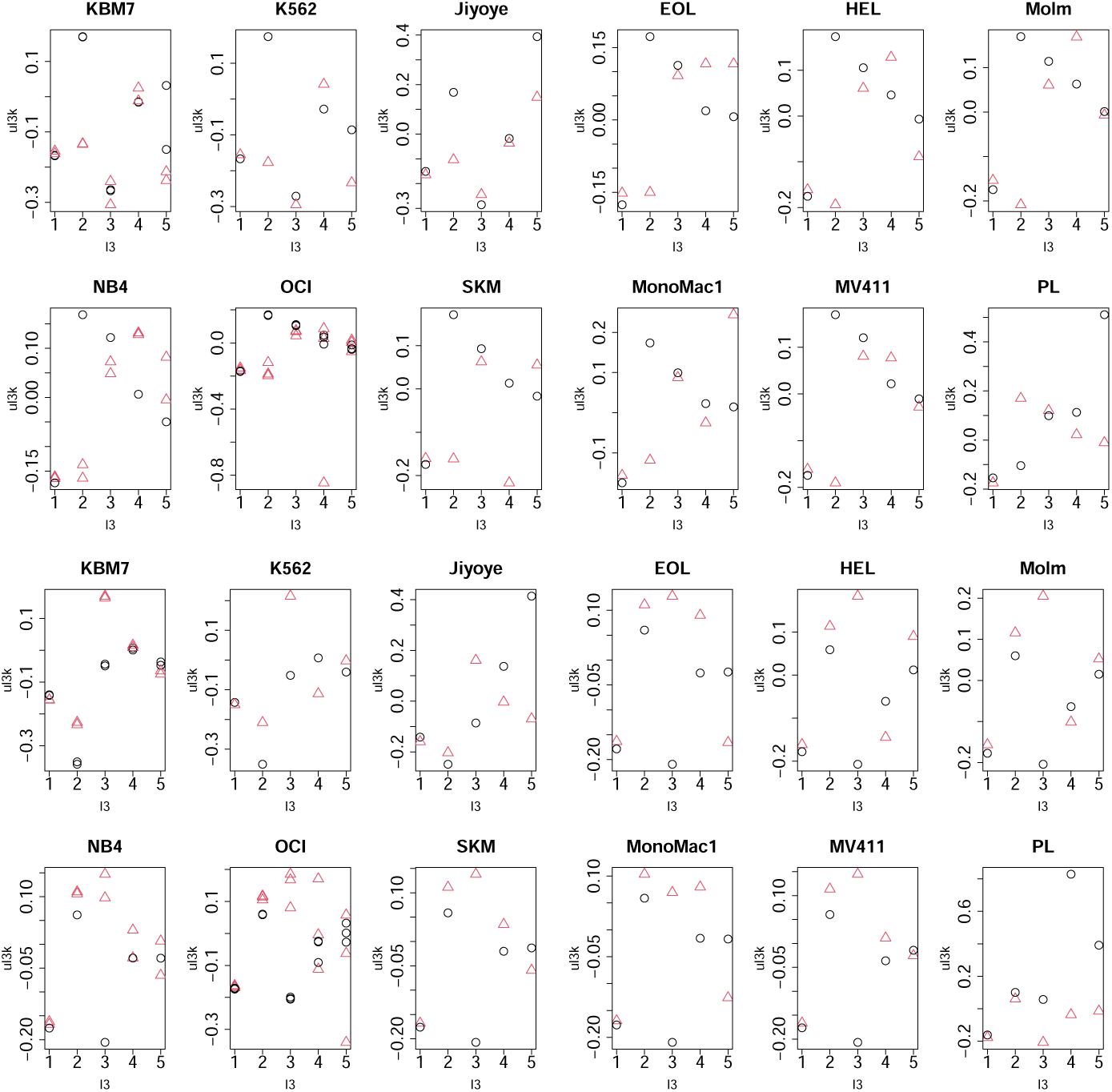
The singular value vectors attributed to cell lines, *u_ℓ_*_3_ *_k_* , for 12 cell lines in the whitehead dataset. The first and second rows: for raw values; the third and fourth rows: for logarithmic values. Red triangles: final and black circles: initial.

#### 2.2.4 GeCKOv2

The above results that a large AUC is associated with the cell-line-attributed singular value vectors that are associated with a large separation between the control and others are considerably reasonable. Nevertheless, the inclusion of controls is not always required for a large AUC, as shown below. As described in the Methods section, the controls are dropped during the screening process (at least three biological replicates) for GeCKOv2 datasets. Thus, the GeCKOv2 dataset can obtain a large AUC despite missing controls. To consider this, we attempted to interpret the cell-line-attributed singular value vectors associated with the gene-attributed singular value vectors associated with a large AUC without dependence on the control. *u*_2*k*_ was selected (upper two panels in Fig. 8) as the cell-line-attributed singular value vector associated with the largest absolute *G*(*ℓ*_1_*ℓ*_2_*ℓ*_3_*ℓ*_4_) with the selected singular value vector attributed genes, *u*_5*i*_, which have large AUC values (Table 1). We downloaded CRISPR efficacies attributed to cell lines from the DepMap database and found that *u*_2*k*_ values were highly correlated with the downloaded CRISPR efficacy attributed to cell lines (lower two panels in Fig. 8).

**Fig. 8.**
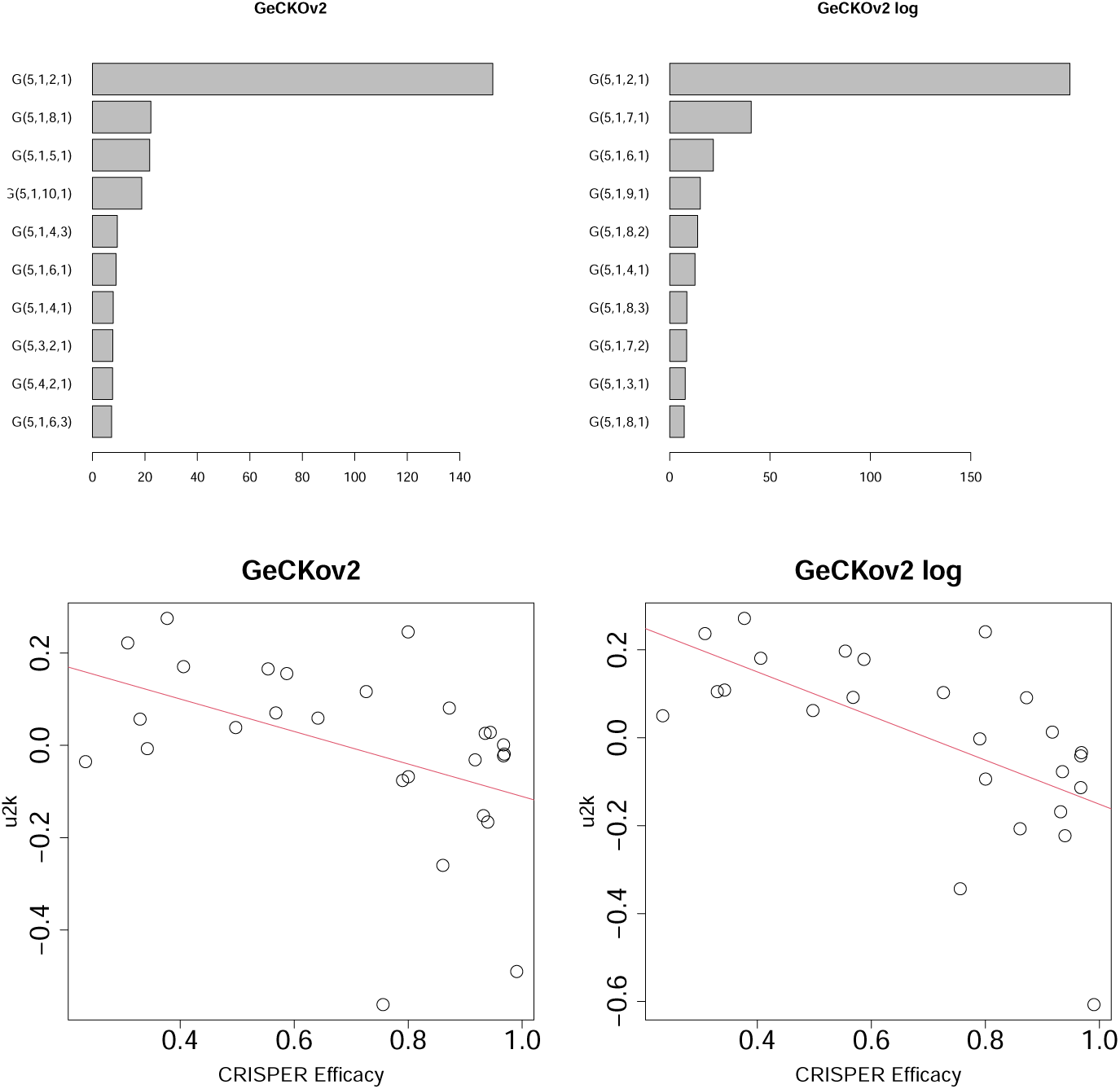
Upper row: The absolute values of *G*(*ℓ*_1_*ℓ*_2_*ℓ*_3_) in the descending order for the GeCKOv2 dataset. *ℓ*_3_ = 2 is selected for raw (left) and logarithmic vales (right). Lower row: Scatter plot between *u*_2*k*_ and CRISPR efficacy. Correlation coefficients are -0.45 (left, raw values) and -0.62 (right, logarithmic values), respectively. Red lines show thez regression lines.

#### 2.2.5 Avana

This section discusses the Avana dataset. *u*_3*j*_ and *u*_2*j*_ were selected as the raw and logarithmic values, respectively (upper two panels in Fig. 9) as cell-line-attributed singular value vectors associated with the largest absolute *G*(*ℓ*_1_*ℓ*_2_*ℓ*_3_*ℓ*_4_) with the selected singular value vectors attributed genes, *u*_5*i*_, that have large AUC values (Table 1). A high AUC is associated with a high correlation between CRISPR efficacy and the cell line-attributed singular value vectors *u*_3*j*_ and *u*_2*j*_ (lower two panels in Fig. 9). Avana dataset also lacks controls.

**Fig. 9.**
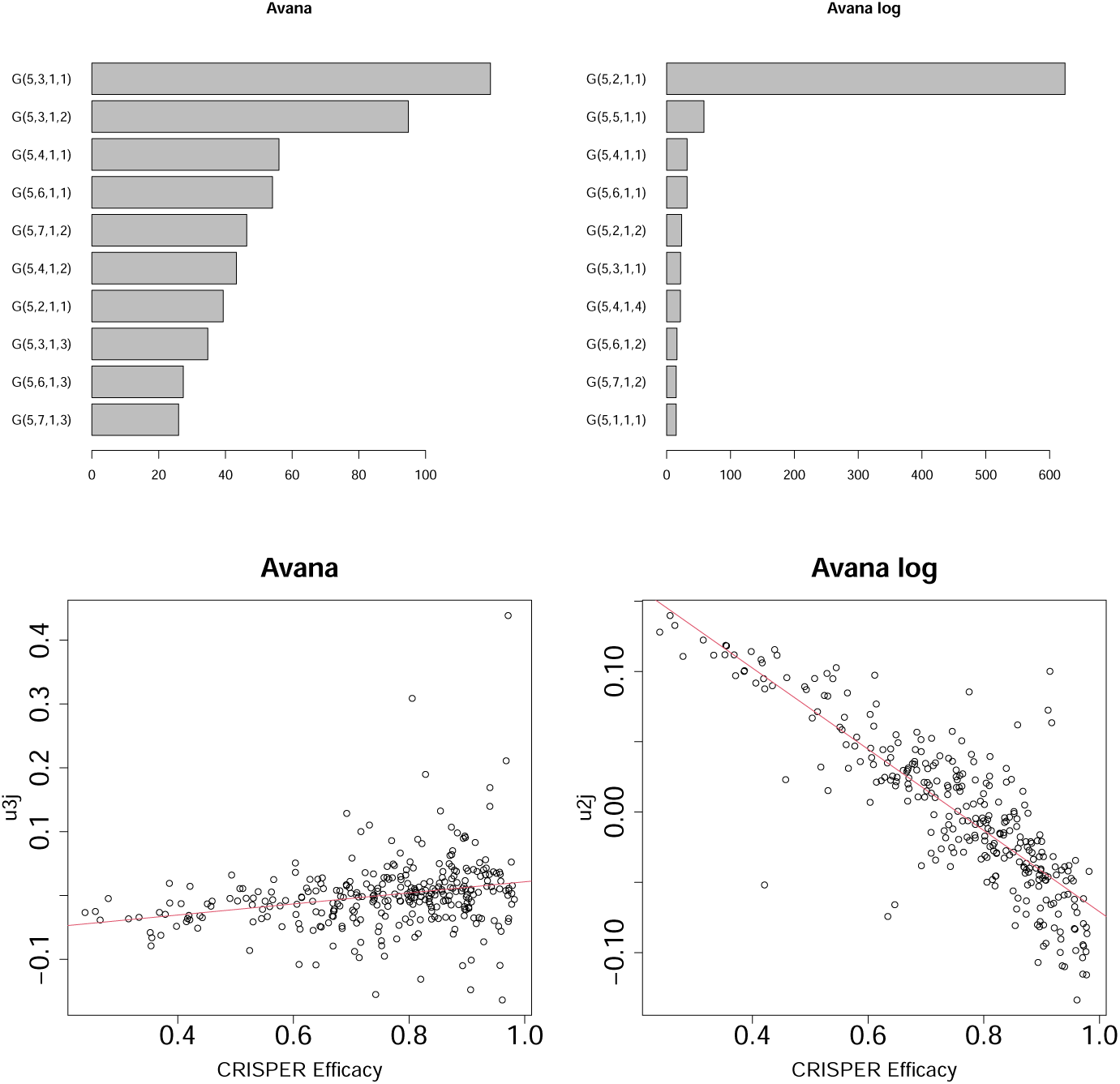
Upper row: The absolute values of *G*(*ℓ*_1_*ℓ*_2_*ℓ*_3_) in descending order for the Avana dataset. *ℓ*_2_ = 3 and *ℓ*_2_ = 2 are selected for raw (left) and logarithmic vales (right), respectively. Lower row: Scatter plot between *u*_3*j*_ (left, raw values) or *u*_2*j*_ (right, logarithmic values) and CRISPR efficacy. Correlation coefficients are 0.25 (left, raw values) and -0.84 (right, logarithmic values), respectively. Red lines show the regression lines.

Thus, TD can achieve a high AUC by identifying cell line-attributed singular value vectors associated with CRISPR efficacy, even in the absence of controls.

## 3 Methods

### 3.1 sgRNA profiles and tensor format

We employed five datasets composed of sgRNA profiles tested in the JACKS paper [9], which were formatted as tensors for downward analysis. All sgRNA profile files were retrieved from the Data.zip available at https://www.doi.org/10.6084/m9.figshare.6002438.

#### 3.1.1 Dataset 1: Yusa V1.0

The file extracted from Data.zip and used for analysis was yusa raw v10.tab. Yusa V1.0 [11, 12] is composed of 16 cell lines and one control. Because the number of replicates associated with individual cell lines varied, 53 profiles were included. Although the number of sgRNAs targeting individual genes also varied, we considered 17,019 genes associated with at least five sgRNAs. When more than five sgRNAs targeted a single gene, five randomly selected sgRNAs were considered. Consequently, we obtained the tensor *x_ijk_*∈ R^17019*×*5*×*53^ which represents the number of *j* sgRNAs that target the *i*th gene of the *k*th profile.

#### 3.1.2 Dataset 2: TKOv1

The file extracted from Data.zip and used for the analysis was tko counts.txt. TKOv1 [13] comprises six cell lines. Because the number of replicates associated with individual cell lines varied, 61 profiles were included. Although the number of sgRNAs targeting individual genes varied, we considered 14,770 genes associated with at least four sgRNAs. When more than four sgRNAs targeted a single gene, four randomly selected sgRNAs were considered. Consequently, we obtained the tensor *x_ijk_* ∈ R^14770*×*4*×*61^ which represents the number of *j* sgRNAs that target the *i*th gene of the *k*th profile. Remarkably, all cell lines were associated with at least one T0 sample (i.e., untreated) within the replicates.

#### 3.1.3 Dataset 3: Whitehead

The file extracted from Data.zip and used for analysis was Wang2015 2017 merged counts.txt. Whitehead [14, 15] comprises 12 cell lines. Because the number of replicates associated with individual cell lines varied, 37 profiles were included. Although the number of sgRNAs that target individual genes also varied, we considered 18,467 genes to be associated with at least 10 sgRNAs. When more than 10 sgRNAs target a single genes, 10 randomly selected sgRNAs were considered. Consequently, we obtained the tensor *x_ijk_* ∈ R^18647*×*10*×*37^ which represents the number of *j* sgRNAs that target the *i*th gene of the *k*th profile. Remarkably, all cell lines were associated with pairs of replicates composed of initials and finals, where the initials were regarded as controls (not treated).

#### 3.1.4 Dataset 4: GeCKOv2

The file extracted from Data.zip and used for the analysis was Achilles raw GeckoV2.tab. GeCKOv2 [16, 17] contains 36 cell lines, including a control. As the number of replicates associated with individual cell lines varied, 129 profiles were included. We considered 29 cell lines associated with at least four replicates (after screening, controls were excluded). Although the number of sgRNAs targeting individual genes also varied, we considered 18,862 genes associated with at least four sgRNAs. When more than four sgRNAs targeted a single gene, four randomly selected sgRNAs were considered. Consequently, we obtained the tensor *x_ijkm_* ∈ R^18862*×*4*×*29*×*4^ which represents the number of *j*th sgRNAs that target the *i*th gene of the *m*th biological replicate of the *k*th cell line.

#### 3.1.5 Dataset 5: Avana

The file extracted from Data.zip and used for the analysis was Avana sgrna raw readcounts matched.csv, and the corresponding gene names were from Avana sgrnamapping.csv. Avana [18] comprised 335 cell lines without a control. Because the number of replicates associated with individual cell lines varied, 808 profiles were included. We used 317 cell lines with at least two replicates. Although the number of sgRNAs targeting individual genes varied, we considered 17,390 genes associated with at least four sgRNAs. When more than four sgRNAs targeted a single gene, four randomly selected sgRNAs were considered. Consequently, we obtained the tensor *x_ijkm_*∈ R^17390*×*317*×*2*×*4^ which represents the number of *m* sgRNAs targeting the *i*th gene of the *k*th biological replicate of the *j*th cell line.

### 3.2 TD

The TD was performed using HOSVD [10]. Tensors, *x_ijk_* ∈ R*^I×J×K^* or *x_ijkm_* ∈ R*^I×J×K×M^* , are decomposed as follows:

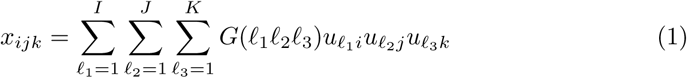

or

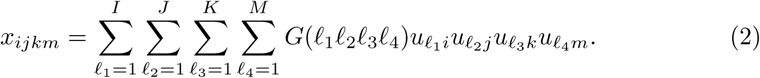

where *G*(*ℓ*_1_*ℓ*_2_*ℓ*_3_) ∈ R*^I×J×K^* or *G*(*ℓ*_1_*ℓ*_2_*ℓ*_3_*ℓ*_4_) ∈ R*^I×J×K×M^* denotes the core tensor representing the contribution of *u_ℓ_*_1 *i*_*u_ℓ_*_2 *j*_*u_ℓ_*_3 *k*_ or *u_ℓ_*_1 *i*_*u_ℓ_*_2 *j*_*u_ℓ_*_3 *k*_*u_ℓ_*_4 *m*_ to *x_ijk_* or *x_ijkm_*, respectively.

Note that the tensors are standardized as follows before being decomposed:

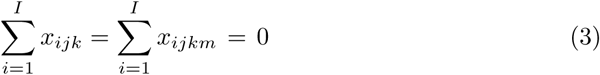

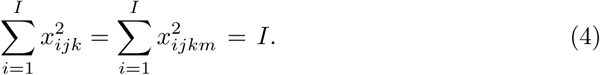

In addition, we consider an alternative version that is logarithmically transformed before standardization, eq. (3) and (4), respectively.

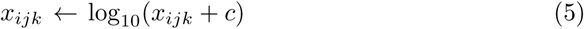

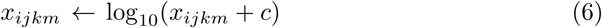

where *c* = 1 or *c* = 2, the smaller of which is employed in individual datasets to satisfy *x_ijk_* + *c >* 0 or *x_ijkm_* + *c >* 0. We also used logarithmically transformed values because various conventional methods employed the logarithmically transformed values.

### 3.3 Retrieval of the lists of essential and non-essential genes

The genes used to evaluate the performance of JACKS and TD were essential and nonessential ones [19]; the numbers of essential and nonessential genes were 360 and 927, respectively. A total of 927 nonessential genes were retrieved from the fifth column of dataset S1. Since there is no explicit list of the 360 essential genes, we used the 823 genes assoicated with “Num Obs” larger than or equal to 3, in the “essential-genes-top12screens” sheet in dataset S4. Here, “Num Obs” is the number of cell lines (of the top 12) in which the gene is classified as essential.

### 3.4 Selection of singular value vectors used

First, we attempted to determine which singular value vectors attributed to genes best discriminated between essential and non-essential genes. Subsequently, we investigated *G* to determine which singular value vectors attributed to the cell lines were mostly related to the selected singular value vectors attributed to cell lines. The selected singular value vectors attributed to the cell lines enabled us to interpret the meaning of the selected singular value vectors attributed to genes that best discriminated between essential and non-essential genes.

### 3.5 Comparison with JACKS

JACKS results were retrieved from JACKS.zip and are available at https://www.doi.org/10.6084/m9.figshare.6002438. Files used are those named as “cell line names” gene pval JACKS results.txt (“cell line names” is “avana,” “gecko2,” “whitehead”, “tko”, or “yusa v10”). Before comparison, *P* -values in datasets are logarithmic-transformed to recover the raw values. We did not use raw values (i.e. files named as “cell line names” gene JACKS results.txt) because logarithmically transformed *P* -values are more robust (i.e., free from the standardization). Because JACKS does not provide a cell line-wide AUC, we computed the mean score of the cell lines in each of the five datasets and used the mean value to compute the AUC.

### 3.6 CRISPR efficacy of cell lines

The CRISPRInferredModelEfficacy.csv file, which included the CRISPR efficiency of the cell lines, was retrieved from DepMap [20]. The “Achilles-Avana-2D” column was used to evaluate the biological reliability of singular value vectors attributed to cell lines (for GeCKOvs and Avana). If the correlation between the CRISPR efficiency of cell lines and the singular value vectors attributed to cell lines is sufficiently large, we can conclude that the singular value vectors attributed to cell lines are biologically reliable.

### 3.7 JACKS standard

In addition to the above procedure to compute AUC of JACKS, we recomputed AUC again by the standard procedure. The critical differences are

- Not P-values but efficiency is used.
- AUC of individual cell lines is computed and averaged.

The R code to perform this, jacks auc.R, is available at GitHub site.

### 3.8 The AUC of Chronos

AUC of Chronos for Avana cell lines is computed from DepMap (25Q3) since DepMap employed Chronos. The R code to perform this, depmap chronos mean auc fixed.R, is available at GitHub site.

## 4 Discussion

First, we emphasize that TD is the first method to integrate multiple sgRNA profiles at the beginning. In the other SOTA introduced in the Introduction, that is, MAGeCK, BAGEL, CERES, STARS, and JACKS, individual sgRNA profiles are analyzed individually, and their outcomes are integrated. However, TD can integrate multiple sgRNA profiles. No separate analytical processes were applied to the individual profiles. This outstanding ability to integrate multiple profiles from the beginning may allow TD to achieve a performance comparable to that of JACKS, even though TD is a simple linear algebra.

Second, it is remarkable that TD, which employs simple linear algebra, can compete with one of SOTA methods, JACKS, which is thought to be highly advanced methods. In particular, for the Avana dataset, which has the largest number of cell lines, the performance of TD was considerably similar to that of JACKS, when we consider not only the AUC value but also the shape of the ROC curve (Fig. 2). Moreover, for Avana, the raw and logarithmic values achieved a similar performance (Fig. 2). This also questions the reliability of taking logarithmic values of the number of sgRNA. If raw (not logarithmic) values perform as well as logarithmic values when attempting to discriminate between essential and nonessential genes, what justifies the usage of logarithmic values? Therefore, it might be better to reconsider the definition of quantitative CRISPR efficiency.

TD also has the advantage of compensating for the distinct efficiencies of individual sgRNAs that target the same genes. Notably, the first singular value vectors attributed to sgRNA are always associated with the largest absolute values of *G* (Table 3). For the Yusa dataset, *u_ℓ_*_2_ *_j_* is attributed to sgRNA. In Fig. 3, *G*(7, 1, 4) and *G*(6, 1, 2) with *ℓ*_2_ = 1 have the largest absolute values. For the TKOv1 dataset, *u_ℓ_*_2_ *_j_* was also attributed to sgRNA. In Fig. 4, *G*(1, 1, 1) and *G*(5, 1, 2) with *ℓ*_2_ = 1 have the largest absolute values. For the whitehead dataset, *u_ℓ_*_2_ *_j_* was again attributed to sgRNA. In Fig. 6, *G*(11, 1, 2), *G*(2, 1, 2), and *G*(12, 1, 3) with *ℓ*_2_ = 1 have the largest absolute values. For the GeCKOv2 datasets, *u_ℓ_*_2_ *_j_* was attributed to sgRNA. In Fig. 8, *G*(5, 1, 2, 1) with *ℓ*_2_ = 1 had the largest absolute values. For the Avana dataset, *u_ℓ_*_4_ *_m_* was attributed to sgRNA. In Fig. 9, *G*(5, 3, 1, 1) and *G*(5, 2, 1, 1) with *ℓ*_4_ = 1 had the largest absolute values. Because the first singular value vectors usually correspond to the simple average, the fact that the first singular value vectors attributed to sgRNA are always employed means that TD is free from the sophisticated consideration of sgRNA efficiency. In other words, TD is a robust method that does not need to consider the differences in CRISPR efficiency between multiple sgRNAs. This might be the reason why TD competes with JACKS, which must consider the difference in sgRNA efficiency.

This also explains why a larger *ℓ*_1_ is often attributed to singular value vectors attributed to genes *u_ℓ_*_1_ *_i_* with a large AUC. Since there are *J* (for datasets other than Avana) and *M* (for Avana dataset) singular value vectors attributed to sgRNAs and *J* or *M* singular value vectors attributed to sgRNAs must represent all dependence upon *j* (for datasets other than Avana) or *m* (for Avana dataset), if the first singular value vectors attributed to sgRNAs represent uniform one (i.e., *u*_1*j*_ or *u*_1*m*_ is independent of *j* or *m*), the next singular value vectors attributed to sgRNAs that represent uniform one must be *j > J* or *m > M* . This results in *u_ℓ_*_1_ *_i_* with larger *ℓ*_1_ are associated with the next singular value vectors attributed to sgRNAs that represent a uniform one because a large AUC is associated with singular value vectors attributed to sgRNAs and is independent of *j* or *m*.

TD also has disadvantages, in that genes and cell lines without sufficiently large sgRNAs or biological replicates are discarded. Although we also have some methods for dealing with cases in which different replicates are associated with [21], we did not employ it for simplicity. In the case of a more sophisticated treatment, we may be forced to consider such cases.

Although we have compared our performances with those of JACKS, as a SOTA, achieved, since JACKS is a bit old (released in 2019), it is better to compare our with more updated ones if possible to confirm the superiority of our method. As one of such updated methods, we employed Chronos [22], which was also used in DepMap. Since the performance of Chronos can be derived from the output in DepMap (25Q3), we have computed AUC with the data from DepMap for Avana that includes the largest number of cells among five cell lines tested (Table 2). Since the performance of Chronos is quite similar to JACKS, it confirms the representativity of JACKS as a SOTA.

We used P-values for JACKS instead of the efficiency and AUC is computed after averaging over cell lines. To confirm that it does not affect the performance of JACKS much, we recomputed AUC for JACKS by following the standard procedure. At first, AUC is computed by the efficiency, not by p-values. Then AUC is averaged over AUC of individual cell lines. Then we have found that this does not change the performance of JACKS much.

## 5 Conclusion

In this study, we applied TD to five datasets comprising sgRNA profiles. TD can integrate sgRNAs and profiles simultaneously and discriminate between essential and non-essential genes as well as JACKS, a type of SOTA. Because TD employs simple linear algebra, it is remarkable that TD can compete with JACKS.

## Supplementary information

## Acknowledgements

Acknowledgements are not compulsory. Where included they should be brief. Grant or contribution numbers may be acknowledged.

Please refer to Journal-level guidance for any specific requirements.

## Declarations

Some journals require declarations to be submitted in a standardised format. Please check the Instructions for Authors of the journal to which you are submitting to see if you need to complete this section. If yes, your manuscript must contain the following sections under the heading ‘Declarations’:

## Ethics approval and consent to participate

Not applicable

## Competing interests

The authors have no conflicts of interest to declare.

## Funding

This project was funded by the Deanship of Scientific Research (DSR) at King Abdulaziz University, Jeddah, Saudi Arabia under grant no. (IPP: 70-611-2025). The authors, therefore, acknowledge with thanks DSR for technical and financial support.

## Data availability

All data analysed in this study were obtained from https://www.doi.org/10.6084/m9.figshare.6002438.

## Code availability

The sample source code is available at https://github.com/tagtag/TDbasedUFE sgRNA.

## Author contribution

Y. H. T. planned the study and performed the analyses. Y.-H.T. and T.T. evaluated the results and wrote and reviewed the manuscript. All the authors have read and agreed to the published version of this manuscript. The conceptualization, data curation, and analysis were performed by Y. H. T.

## Notes

### Competing Interest Statement

The authors have declared no competing interest.

### Summary of Updates

Two sections were added to the end of Methods and two paragraphs were added to the end of Discussion to address reviewers' comments.

## References

[1] Shalem, O., Sanjana, N.E., Zhang, F.: High-throughput functional genomics using CRISPR–Cas9. Nature Reviews Genetics 16(5), 299–311 (2015) 10.1038/nrg3899

[2] Li, W., Xu, H., Sizemore, T., Love, M.I., Zhang, F., Meyer, C.A., Liu, J.S., Brown, M., Liu, X.S.: MAGeCK enables robust identification of essential genes from genome-scale CRISPR/Cas9 knockout screens. Genome Biology 15(12), 554 (2014) 10.1186/s13059-014-0554-4

[3] Li, W., Koster, J., Xu, H., Chen, C.-H., Sizemore, T., Liu, J.S., Liu, X.S.: Quality control, modeling, and visualization of CRISPR screens with MAGeCK-VISPR. Genome Biology 16(1), 271 (2015) 10.1186/s13059-015-0843-6

[4] Hart, T., Moffat, J.: BAGEL: a computational framework for identifying essential genes from pooled library screens. BMC Bioinformatics 17(1), 164 (2016) 10.1186/s12859-016-1015-8

[5] Doench, J.G., Fusi, N., Sullender, M., Hegde, M., Vaimberg, E.W., Donovan, K.F., Smith, I., Tothova, Z., Wilen, C., Orchard, R., Virgin, H.W., Listgarten, J., Root, D.E.: Optimized sgRNA design to maximize activity and minimize off-target effects of CRISPR-Cas9. Nature Biotechnology 34(2), 184–191 (2016) 10.1038/nbt.3437 . Epub 2016 Jan 18

[6] Hsu, P.D., Scott, D.A., Weinstein, J.A., Ran, F.A., Konermann, S., Agarwala, V., Li, Y., Fine, E.J., Wu, X., Shalem, O., Cradick, T.J., Marraffini, L.A., Bao, G., Zhang, F.: Dna targeting specificity of rna-guided Cas9 nucleases. Nature Biotechnology 31(9), 827–832 (2013) 10.1038/nbt.2647 . Epub 2013 Jul 21

[7] Meyers, R.M., Bryan, J.G., McFarland, J.M., Weir, B.A., Sizemore, A.E., Xu, H., Dharia, N.V., Montgomery, P.G., Cowley, G.S., Pantel, R., Brügmann, A.L.M., Kim, E., Vilfort, Z., Tsherniak, A., Vazquez, F., Adstamongkonkul, P., Mermel, C.H., Doench, J.G., Root, D.E., Ching, T.H.N., Boehm, J.S., Hahn, W.C.: Computational correction of copy number effect in CRISPR–Cas9 screens. Nature Genetics 49(12), 1779–1784 (2017) 10.1038/ng.3980

[8] Doench, J.G., Hartenian, E., Brown, K.R., Root, D.E., Lander, E.S.: Rational design of highly active sgRNAs for CRISPR-Cas9-mediated gene inactivation. Nature Biotechnology 32(12), 1262–1267 (2014) 10.1038/nbt.3026

[9] Allen, F., Behan, F., Khodak, A., Iorio, F., Yusa, K., Garnett, M.J.: JACKS: joint analysis of CRISPR/Cas9 knockout screens. Genome Research 29(3), 464–471 (2019) 10.1101/gr.238710.118

[10] Taguchi, Y.-h.: Unsupervised Feature Extraction Applied to Bioinformatics: A PCA Based and TD Based Approach, 2nd edn. Unsupervised and Semi-Supervised Learning. Springer, Switzland (2024)

[11] Tzelepis, K., Koike-Yusa, H., De Braekeleer, E., Li, Y., Metzakopian, E., Dovey, O.M., Mupo, A., Grinkevich, V., Li, M., Mazan, M., Gozdecka, M., Ohnishi, S., Cooper, J., Patel, M., McKerrell, T., Chen, B., Domingues, A.F., Gallipoli, P., Teichmann, S., Ponstingl, H., McDermott, U., Saez-Rodriguez, J., Huntly, B.J.P., Iorio, F., Pina, C., Vassiliou, G.S., Yusa, K.: A crispr dropout screen identifies genetic vulnerabilities and therapeutic targets in acute myeloid leukemia. Cell Reports 17(4), 1193–1205 (2016) 10.1016/j.celrep.2016.09.079. doi: 10.1016/j.celrep.2016.09.079

[12] Iorio, F., Behan, F.M., Gonçalves, E., Bhosle, S.G., Chen, E., Shepherd, R., Beaver, C., Ansari, R., Pooley, R., Wilkinson, P., Harper, S., Butler, A.P., Stronach, E.A., Saez-Rodriguez, J., Yusa, K., Garnett, M.J.: Unsupervised correction of gene-independent cell responses to CRISPR-Cas9 targeting. BMC Genomics 19(1), 604 (2018) 10.1186/s12864-018-4989-y

[13] Hart, T., Chandrashekhar, M., Aregger, M., Steinhart, Z., Brown, K.R., MacLeod, G., Mis, M., Zimmermann, M., Fradet-Turcotte, A., Sun, S., Mero, P., Dirks, P., Sidhu, S., Roth, F.P., Rissland, O.S., Durocher, D., Angers, S., Moffat, J.: High-resolution crispr screens reveal fitness genes and genotype-specific cancer liabilities. Cell 163(6), 1515–1526 (2015) 10.1016/j.cell.2015.11.015 . doi: 10.1016/j.cell.2015.11.015

[14] Wang, T., Birsoy, K., Hughes, N.W., Krupczak, K.M., Post, Y., Wei, J.J., Lander, E.S., Sabatini, D.M.: Identification and characterization of essential genes in the human genome. Science 350(6264), 1096–1101 (2015) 10.1126/science.aac7041 https://www.science.org/doi/pdf/10.1126/science.aac7041

[15] Wang, T., Yu, H., Hughes, N.W., Liu, B., Kendirli, A., Klein, K., Chen, W.W., Lander, E.S., Sabatini, D.M.: Gene essentiality profiling reveals gene networks and synthetic lethal interactions with oncogenic ras. Cell 168(5), 890–90315 (2017) 10.1016/j.cell.2017.01.013 . doi: 10.1016/j.cell.2017.01.013

[16] Sanjana, N.E., Shalem, O., Zhang, F.: Improved vectors and genome-wide libraries for crispr screening. Nature Methods 11(8), 783–784 (2014) 10.1038/nmeth.3047

[17] Aguirre, A.J., Meyers, R.M., Weir, B.A., Vazquez, F., Zhang, C.-Z., Ben-David, U., Cook, A., Ha, G., Harrington, W.F., Doshi, M.B., Kost-Alimova, M., Gill, S., Xu, H., Ali, L.D., Jiang, G., Pantel, S., Lee, Y., Goodale, A., Cherniack, A.D., Oh, C., Kryukov, G., Cowley, G.S., Garraway, L.A., Stegmaier, K., Roberts, C.W., Golub, T.R., Meyerson, M., Root, D.E., Tsherniak, A., Hahn, W.C.: Genomic copy number dictates a gene-independent cell response to crispr/cas9 targeting. Cancer Discovery 6(8), 914–929 (2016) 10.1158/2159-8290.CD-16-0154 . This article is highlighted in the In This Issue feature, p. 803

[18] Meyers, R.M., Bryan, J.G., McFarland, J.M., Weir, B.A., Sizemore, A.E., Xu, H., Dharia, N.V., Montgomery, P.G., Cowley, G.S., Pantel, S., Goodale, A., Lee, Y., Ali, L.D., Jiang, G., Lubonja, R., Harrington, W.F., Strickland, M., Wu, T., Hawes, D.C., Zhivich, V.A., Wyatt, M.R., Kalani, Z., Chang, J.J., Okamoto, M., Stegmaier, K., Golub, T.R., Boehm, J.S., Vazquez, F., Root, D.E., Hahn, W.C., Tsherniak, A.: Computational correction of copy number effect improves specificity of crispr–cas9 essentiality screens in cancer cells. Nature Genetics 49(12), 1779–1784 (2017) 10.1038/ng.3984

[19] Hart, T., Brown, K.R., Sircoulomb, F., Rottapel, R., Mof-fat, J.: Measuring error rates in genomic perturbation screens: gold standards for human functional genomics. Molecular Systems Biology 10(7), 733 (2014) 10.15252/msb.20145216 https://www.embopress.org/doi/pdf/10.15252/msb.20145216

[20] Tsherniak, A., al.: Defining a cancer dependency map. Cell 170(3), 564–576 (2017) 10.1016/j.cell.2017.06.010

[21] Taguchi, Y.-h., Turki, T.: A tensor decomposition-based integrated analysis applicable to multiple gene expression profiles without sample matching. Scientific Reports 12(1), 21242 (2022) 10.1038/s41598-022-25524-4

[22] Dempster, J.M., Boyle, I., Vazquez, F., Root, D.E., Boehm, J.S., Hahn, W.C., Tsherniak, A., McFarland, J.M.: Chronos: a cell population dynamics model of CRISPR experiments that improves inference of gene fitness effects. Genome Biology 22(1), 343 (2021) 10.1186/s13059-021-02540-7

